# A methodology for co-simulation-based optimization of biofabrication protocols

**DOI:** 10.1101/2022.01.28.478198

**Authors:** Leonardo Giannantoni, Roberta Bardini, Stefano Di Carlo

## Abstract

Biofabrication processes are complex and often unsatisfactory. Trial-and-error methods are costly and yield only incremental innovation, starting from sub-optimal and poorly represented existing processes. Although computational techniques might support efficient process design to find optimal process configurations, intelligent computational approaches must comprise biological complexity to provide meaningful insights. This paper proposes a novel co-simulation-based optimization methodology for the systematic design of protocols for cell culture and biofabrication. The proposed strategy integrates evolutionary computation and simulation for efficient design space exploration and assessment of candidate protocols. A generic library supports the modular and flexible composition of multiscale and multidomain co-simulation scenarios. The feasibility of the presented approach was demonstrated in the automatic generation of protocols for the biofabrication of an epithelial cell monolayer. The results are twofold. First, the prototype co-simulation library helps build flexible, loosely coupled simulation scenarios. Second, the in-silico experimentation on the use case shows that the proposed approach is a viable first step towards standard and automated design in biofabrication.

## 1 Introduction

Biofabrication is “*the automated generation of biologically functional products with the structural organization from living cells, bioactive molecules, biomaterials, cell aggregates such as micro-tissues, or hybrid cell-material constructs, through Bioprinting or Bioassembly and subsequent tissue maturation processes*” [8]. Biofabrication in Tissue Engineering and Regenerative Medicine (TERM) has the potential to disrupt clinical and pharmacological research [19]. Yet, biofabrication of complex and large tissues and organs is still out of reach.

Biofabrication processes are highly complex biologically and technologically. Biofabrication requires the application of specific protocols representing the dynamic configuration of relevant process control parameters, emphasizing the values they assume in space and time. However, the large number of critical parameters implies a vast design space, whose exploration is prohibitive and impairs the results obtainable by common *in vitro* trial-and-error experiments [4]. These include brute-force experimental campaigns and One Factor at A Time (OFAT) strategies exploring ranges of relevant system parameters one at a time while holding the others constant [7]. This approach is expensive in terms of time and resources. Also, it overlooks inter-dependencies among system variables, which impedes linking experimental results with process designs controlling multiple variables at a time. This can result in sub-optimal processes [26].

Automation [10] and digitalization [9, 26] make trial-and-error approaches more efficient, reducing operator-dependency and human errors, thus supporting process tracking and control. This dramatically increases the yield of *in vitro* experimental campaigns, allowing a more significant number of experiments, thus a broader exploration of the design space. Yet, making the execution of experiments more efficient does not affect the underlying trial-and-error paradigm. *In silico* Design Space Exploration (DSE) approaches can instead support research design and optimization to maximize information extraction, and process improvement efficiency [12].

This work presents the first step towards optimization via simulation (OvS) for generating optimal biofabrication protocols for defined target products. In particular, the proposed framework follows the model-based simulation-optimization paradigm in which DSE and simulation modules are tightly integrated. The DSE selects the solutions which need to be evaluated by simulation [1]. The proposed method exploits heuristic DSE based on Genetic Algorithms (GA) to increase computational feasibility and combines it with a co-simulation environment relying on white-box simulation models to maximize expressivity and explainability. The original contribution of this paper also includes a library of generic components supporting the modular and flexible composition of co-simulation scenarios. Therefore, the co-simulation can easily combine different models, including the target biological entities (e.g., cells), the biofabrication environment, and the possible stimuli delivered during the biofabrication process. The entire framework is presented, resorting to a selected use case to generate optimal protocols for cultivating two-dimensional epithelial sheets with specific shapes relying on a model including both intracellular and extracellular processes. Experimental results show the capability of the proposed approach and identify a set of significant challenges to stimulate further research in this field.

## 2 Background

Model-based simulation-optimization techniques are part of the broader field of computational process design for biofabrication. Several computational methodologies for biofabrication process design exist in the state-of-the-art. Design-of-Experiments (DoE) [23, 26] supports strategic and effective research design by enabling efficient, systematic exploration and exploitation of complex design spaces [7, 14]. A variety of DoE approaches exist [11], and they prove adequate to tackle multi-factorial problems in the optimization of directed cell differentiation [3, 18], and tissue engineering scaffolds [25]. DoE can be combined with Machine Learning (ML) and Artificial Neural Networks (ANN) to improve the accuracy of the bioprocess model [23].

Yet, ML and ANN provide black-box models of the system. Comprehensive modeling of biological complexity is critical for developing computational approaches for biofabrication [5]. To support informed decisions in process design, the ideal model of biofabrication must be accurate, predictive, interpretable and able to analyze process dynamics. Computer simulations provide white-box models of the process, a powerful tool for analyzing complex systems, and particularly their trajectories under different conditions [1].

Simulation and optimization can work together. In optimization via simulation (OvS) methods, optimization can leverage simulations to explore the process design space, and DoE can support the design of simulations campaigns [11]. OvS leverages the simulation model of a physical process to explore its dynamic behavior after specific stimuli, where the parameter values are systematically varied to find the most performing combination towards a target objective [2]. OvS includes model-based and metamodel-based approaches [1]. In model-based OvS, the optimization engine selects the solutions evaluated by simulation. Modelbased approaches combine the accuracy and interpretability of simulations with the systematic exploration of the process design space provided by optimization. This fulfills the requirements of biofabrication process optimization, yet it poses strong limitations in terms of computational feasibility when the modeled system is complex. A strategy to reduce the computational complexity of OvS is to include a metamodel that estimates input-output relations of the simulation model to significantly reduce the computational time at the cost of accuracy [17] and a partial fallback to black-box modeling. Also, in this case, white-box is preferable to black-box modeling since interpretability and explainability of OvS results build their relevance for the design of an actual biofabrication process. Heuristic methods allow us to find an approximate solution faster than full-space search methods by trading accuracy and completeness for speed while maintaining the simulated model intact. Among them, the Genetic Algorithm (GA) mimics biological evolutionary dynamics where solutions in the design space undergo a process similar to natural selection [15].

## 3 Methods

Figure 1 summarizes the main architecture of the presented framework, including a Design Space Exploration (DSE) engine for the generation of biofabrication protocols and a simulation engine for testing them. The framework receives the high-level specification of the target product and iteratively computes a biofabrication protocol optimized to grow it. The DSE assembles potential biofabrication protocols and feeds them to simulation instances. Simulation results are compared against the specifications of the target product used to rank the corresponding protocols and generate new ones at the next iteration. This procedure continues until an optimal protocol is produced, a predetermined number of iterations is reached, or the protocol performance stalls for a given number of iterations. To help the reader, the paper introduces the proposed framework using a running use case focusing on the fabrication of a human epithelial cells monolayer with selected shapes.

**Fig. 1.**
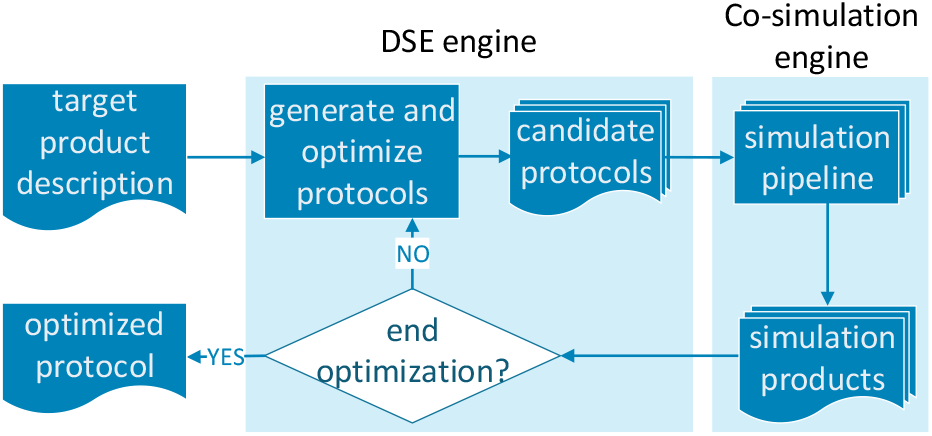
A high-level representation of the simulation-optimization pipeline. Given a target product, the DSE engine generates a pool of candidate protocols to be simulated. The products obtained by simulation are compared to the desired target. The previous steps are iterated until an optimal protocol is found.

### 3.1 Use case description

As a proof of concept, this work presents the generation of protocols for fabricating human epithelial sheets. To this end, the proposed use case includes a computational model of a population of epithelial cells, modeling intracellular and extracellular processes.

The intracellular model is a Boolean Network (BN) based on a published and well-documented work synthesizing epithelial cells behavior (i.e., survival, proliferation, and apoptosis) in response to a combination of cues [22]. These include environmental factors such as cell density, extracellular matrix stiffness, and growth factor signaling. The high abstraction level of this model allows for low computational complexity and easy integration of new knowledge. A graphical representation of the used Boolean network is available in Fig. 3 of the above paper.

The extracellular model describes interactions among cells and between cells and the environment. It models a discrete 3D grid supporting cells evolving on an extracellular matrix (ECM) surface and interacting with neighboring cells and environmental stimuli.

Biofabrication of a target product can be guided in this model by administering growth factors (GF) at a given 3D coordinate, i.e., molecules that stimulate cell proliferation, and by exposing it to TNF-related apoptosis-inducing ligand (TRAIL), a protein inducing cell death by apoptosis [24]. The biofabrication process can also control the deposition of cells in the culturing environment.

The two models are coupled and interact through specific inputs. For instance, the intracellular model includes a `CellDensity_High` node, which is used by the Boolean equation to determine the value for the `Replication` node. If, according to the extracellular model, a cell ends up in a very dense area, the cell is informed by setting its `CellDensity_High` to `True`. This, in turn, affects the cell’s ability to replicate, thus simulating the inhibition of proliferation by contact inhibition.

### 3.2 Co-Simulation engine

The co-simulation engine interacts with the DSE engine to simulate and evaluate the candidate biofabrication protocols. It sets up and evolves the biological system for a predetermined number of simulation steps, administering stimuli according to the protocol under test. As described in subsection 3.1, biofabrication requires the co-simulation of intertwined aspects, each based on a different formalism. Therefore, different simulators must be connected through an interface to exchange data and handle different scales and domains. Several freely-available libraries were tested to ease such implementation (e.g., mosaik [21]). However, they cannot dynamically change the topology of the simulators required to handle the intrinsic dynamical nature of biological systems.

To overcome this limitation, a prototypal co-simulation framework was developed. It is a Python library of generic components that can set up loosely-coupled co-simulation scenarios, either standalone or associated with a DSE engine. It supports multiscale and multidomain systems and provides mechanisms for transparent distributed execution and third-party software encapsulation.

Figure 2(a) depicts the overall architecture of the simulation framework. A Simulation encapsulates a `Pipeline` of `Simulators`, each executing an arbitrary number of `Model` entities. This design provides common interfaces to ensure interoperability between multiscale and multidomain simulators, either custom or pre-existing, that can be transparently instantiated on local or remote machines relying on the Pyro4 library [13]. Flexible composition and clear separation are coupled with an ad-hoc loose-coupling mechanism (i.e., it does not involve an orchestrator) for information exchange using a shared `Event Dictionary` collecting and relaying all the events in the simulation pipeline. The high degree of modularity and integration does not enforce consistent conceptual interrelations. Therefore, the data exchanged between the simulators might need suitable translation layers provided by intermediate simulators. With this architecture, a simulation scenario can be easily set up through a single file listing the simulators, both local and remote, and their configurations (Listing 1.1).

**Listing 1.1.**
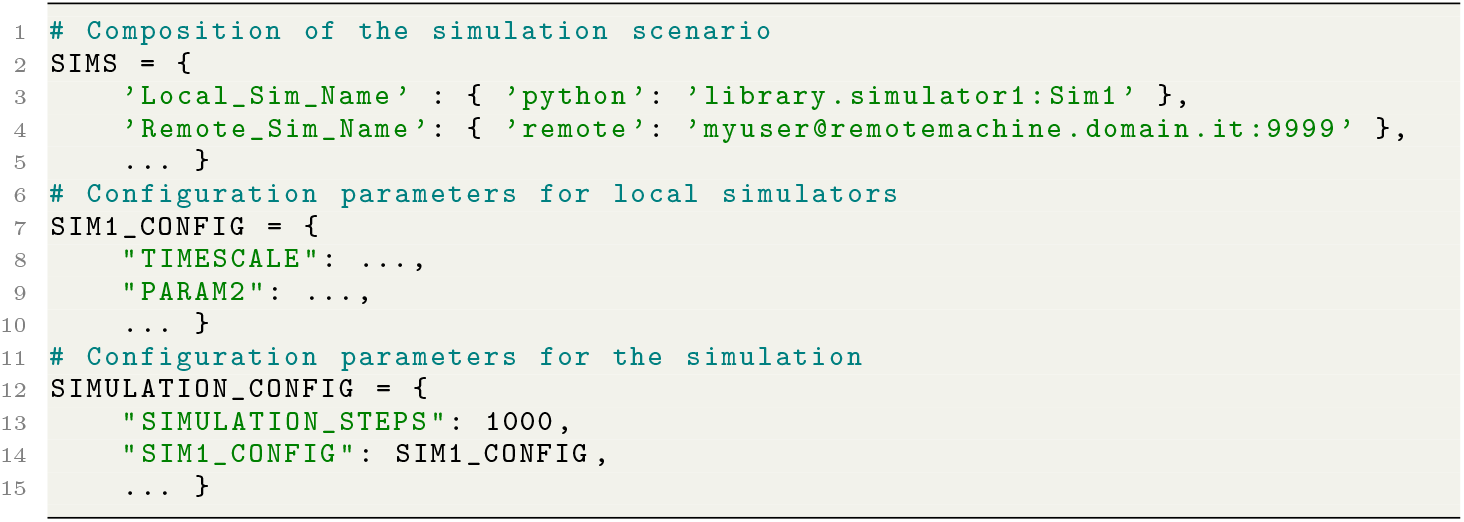
Configuration example for the simulation engine.

**Fig. 2.**
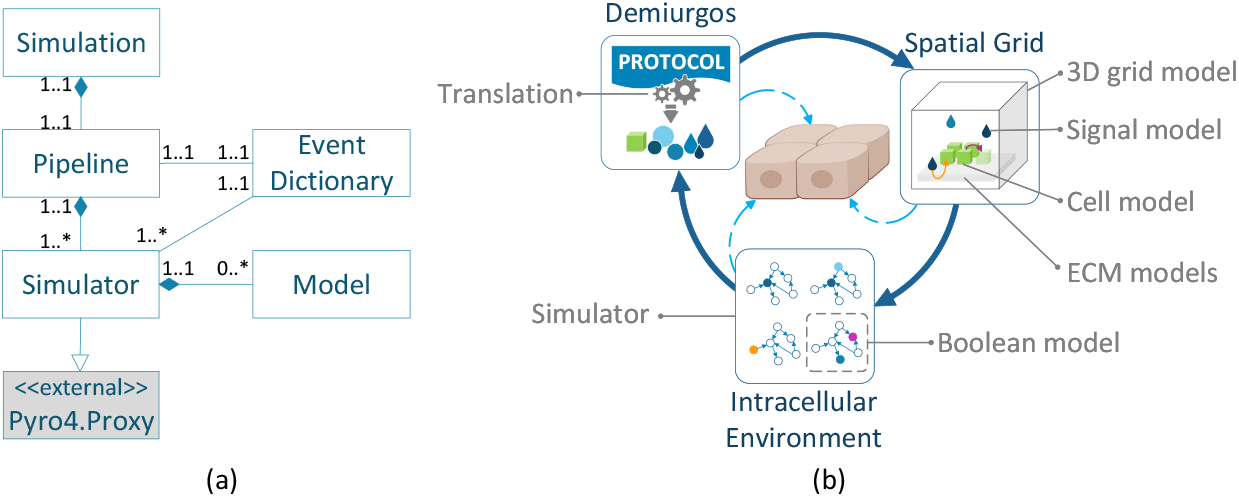
Co-simulation framework. (a) UML diagram of the simulation framework. (b) Co-simulation scenario for the modeled use case.

The considered use case is implemented by the co-simulation setup illustrated in Figure 2(b), supported by the above-described library. Two separate discrete-event simulators, Spatial Grid and Intracellular Environment are dedicated to simulating a discrete 3D grid model with object instances (cells, ECM, signals) and BN models of cells. A third simulator (Demiurgos) translates and administers the protocol commands to the appropriate simulator. Demiurgos, like its Platonic entity namesake, is the means through which protocols manifest in and influence the simulation universe. It acts as a purely functional layer (i.e., it does not simulate any entity) by translating and delivering the culturing protocol under simulation.

The Spatial Grid instantiates cells, ECM, and signal objects, as instructed by Demiurgos, and manages the assignment of unique universal identifiers (UUIDs) to cells. It also mimics the diffusion of GFs and TRAIL in the culturing space with a simplified algorithm. At each simulation step, each signal is displaced according to its drift attribute, which decays over time. Signals are removed from the simulation when their keepalive counter reaches zero. With a random probability, each signal reaching the same coordinate of a cell is tagged to be *consumed* by it and removed from the simulation. If a cell is in an area with a high density of neighboring cells, Spatial Grid prevents replication and communicates to the cell that it is in a high-density area. Such information, in turn, might affect that cell’s behavior in the Intracellular Environment (see subsection 3.1). The Intracellular Environment is informed about a newly issued cell’s UUID if the cell can replicate. The Intracellular Environment manages Boolean network entities modeled after [22] and relies on the PyBoolNet library [16]. It spawns new entities with the UUID provided by Spatial Grid. It feeds them a Boolean input modified by both the protocol and the events coming from the extracellular environment, thus informing them about the cell density of their surroundings, the quality of their supporting ECM, and the availability of nutrients and other signals. Suppose a cell entity enters an apoptotic state. In that case, Intracellular Environment removes it and broadcasts its UUID so that the other simulators in the pipeline (in this case, Spatial Grid) remove the corresponding model too.

### 3.3 Design Space Exploration engine

A biofabrication protocol is a list of signals organized in time and space to guide a specific biological product synthesis. Optimizing such arbitrary long lists of instructions is a complicated combinatorial problem. Therefore, exhaustive exploration is not an option. Our DSE component employs a Genetic Algorithm (GA), an evolutionary computation metaheuristic, to generate populations of candidate solutions or individuals (i.e., the protocols). In this context, a protocol is an individual characterized by a genome whose genes are the signals composing the protocol. Candidate individuals are ranked based on a fitness function and mutated to evolve the population at each new generation. The proposed DSE engine is built on top of the μGP (microGP) library, a tool tailored to problems whose solutions can be expressed similarly to assembly programs [20].

The individuals defined for the proposed use case scenario are organized in two sections (Listing 1.2). The placing section lists the 3D coordinates of the cells to be deposited at the beginning of the biofabrication process. The signal section lists nutrients and environmental stimuli organized in space and time.

**Listing 1.2.**
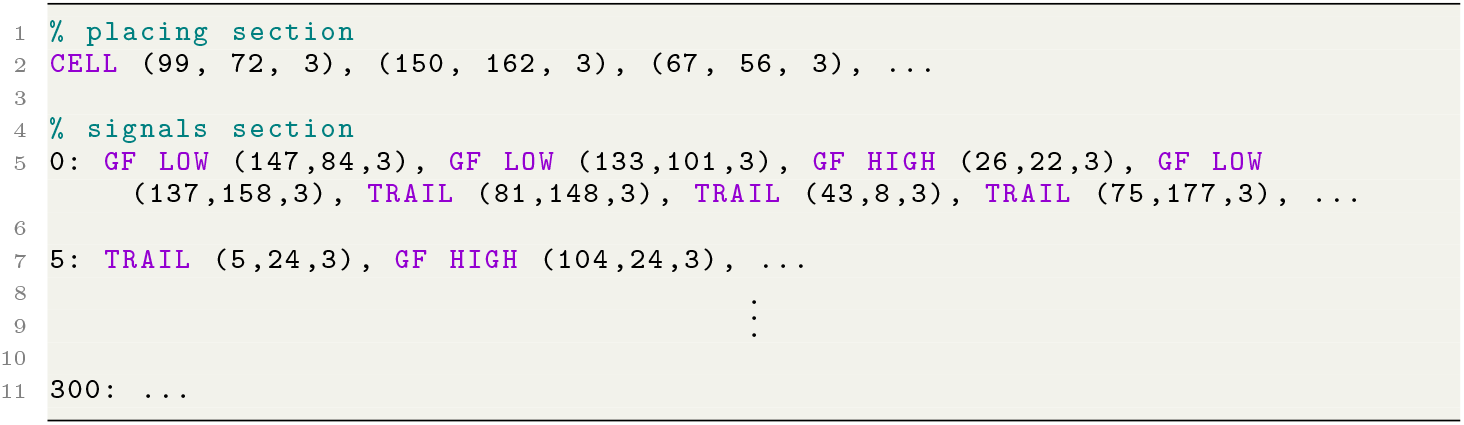
Sample protocol built by the DSE engine.

The *language* used to build the protocols for the epithelial cell model includes `CELL`, `GF LOW/HIGH`, and `TRAIL` macros, each followed by 3D coordinates. They derive from the inputs (cells deposition and exposure to stimuli) specific to the model described in subsection 3.1. This abstract and compact representation minimizes the resources required to compute and store the individuals that in μGP are encoded as directed multigraphs constrained by user-defined rules [20].

The DSE engine requires the specification of a target product, i.e., the biological construct obtained at the end of the simulation and a timescale. In the example provided in Listing 1.3, a `STRIPES` target composed of two parallel planar stripes is described using the sum of two `CUBOID` primitives from our `geometry` library. An optional bounding box can provide cues to the DSE engine for the assembly of new individuals. The timescale allows tuning the granularity of the protocol for the simulated system. For instance, if the protocol step is set to 5, the signal section of the protocol uses a *simulationsteps/*5 length, and Demiurgos issues one protocol instruction every five simulation steps.

The DSE engine uses three genetic operators to evolve the population of candidate protocols.When creating a new offspring, each operator is applied with an initial probability equal to the *strength* parameter *α*, which in our setup is equal to 0.9 (*α* ∈ [0…1]). Strength is a self-adapting parameter. μGP increases it when an operator shows a high success rate (i.e., the mutated individuals’ fitness improved compared to its parents) and decreases it otherwise. `singleParameterAlterationMutation` chooses a new random value for one parameter of an individual. For instance, it might alter the *x* coordinate for the deposition of a signal at protocol step *n*. `onePointCrossover` generates two offspring individuals from two parent individuals by recombining them over a single cut point. For instance, given two 100-step protocols, it might cut them at step 20 and swap the parts containing steps 21 to 100. `twoPointCrossover` operates similarly. It chooses two cut points and swaps the middle portion.

The protocols’ fitness is assessed by comparing the product obtained by simulation with the target product. For the presented application, the fitness is represented using two values:

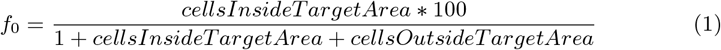

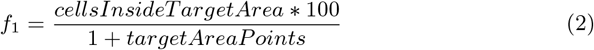

Equation 1 (*f*_0_) expresses the precision, i.e., the fraction of the biofabricated product that matches the target. Equation 2 (*f*_1_) expresses the coverage, i.e., how much of the target product has been obtained. *targetAreaPoints* is defined as the number of integer 3D coordinates included in the target shape. For instance, given a square target covering a *m*×*n* area (i.e., *m×n targetAreaPoints*), a fitness *f* = [84.1, 29.2] means that 29.2% of the desired product has been obtained (i.e., it covers 29.2% of the *m* × *n* area), and 84.1% of the material is where expected (inside the target area). That is, the remaining 15.9 % of cells is misplaced (outside the target area).

μGP provides two methods to evaluate a multi-parameter fitness functions, `Enhanced` and `MultiObjective`. The `Enhanced` method attributes decreasing importance to the *f_i_* parameters. The `MultiObjective` method attributes the same weight to *f*_0_ and *f*_1_, thus leading to the choice of the best individuals based on the joint evaluation of the two parameters. The best individual, in this case, is chosen among those dominating the individuals belonging to the Pareto frontier of the previous generation.

The proposed implementation of the DSE engine employs the `MultiObjective` method, as it is tailored to multi-objective optimization problems. That is, those requiring trade-offs between multiple and potentially conflicting objectives. Cell proliferation is helpful for coverage for the presented use case but must be restrained to avoid abnormal growth. Simultaneously, precision (summarizing a proliferation process under control and limited to a well-defined area) should not prevent obtaining the target product in the desired quantity and shape.

**Listing 1.3.**
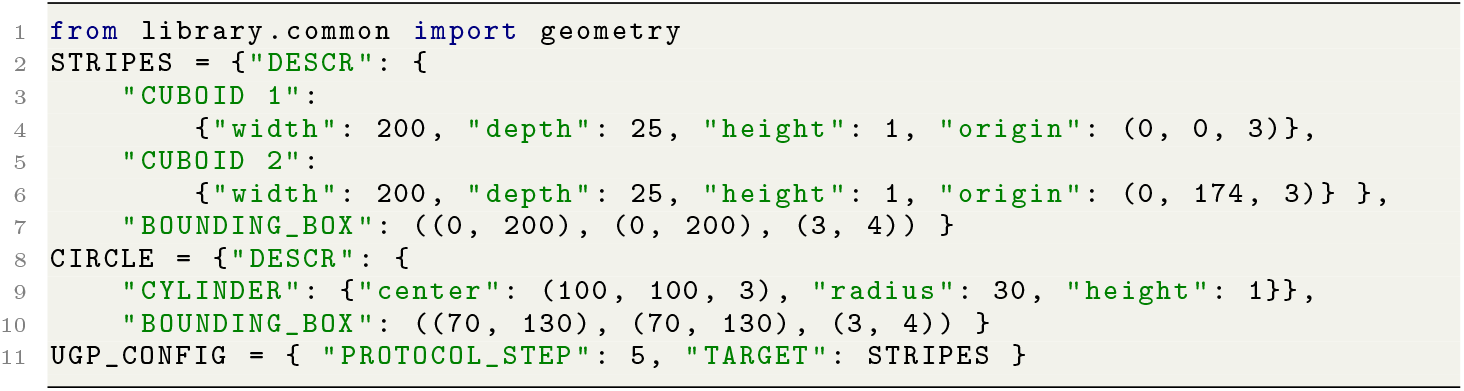
Configuration example for the DSE engine.

## 4 Results

This section presents the validation strategy employed to demonstrate the functioning of the proposed approach.

### 4.1 Experimental setup

The experimental setup starts with the definition of a target product, that is, an epithelial cells monolayer covering half the ECM surface (Figure 3). The culturing environment simulated by the Spatial Grid is a 200 × 200 × 200 cube, with its base covered by a 200 × 200 × 3 layer of ECM entities. The target product is then a 200 × 100 × 1 rectangle lying on the ECM layer. At the beginning of each simulation, the Intracellular Environment sets all new cells in the proliferative state defined in [22]. Demiurgos issues one protocol instruction every five simulation steps. The co-simulation engine evolves the system for 1,500 simulation steps per simulation, stopping in advance if all cells die.

**Fig. 3.**
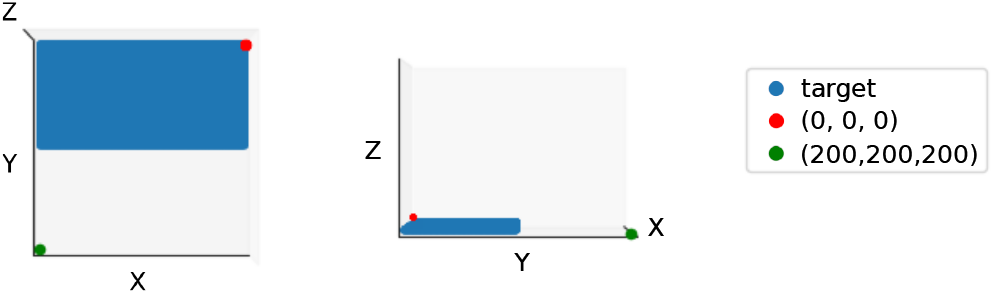
Target product. Left: top view, right: lateral view. The green and red dots help figure out the orientation.

As for this experiment, the placing section of the protocols lists 1 to 15 (*average* = 8, *sigma* = 2) coordinates for cells deposition. The signals section contains 300 (1500/5) instructions, and each instruction contains 0 to 50 (*average* = 15, *sigma* = 5) occurrences of macros (`GF HIGH`, `GF LOW`, and `TRAIL`, as detailed in subsection 3.3). The population evolved by μGP is of *MultiObjective* type, it has an initial size *ν* = 10 and a maximum size *μ* = 10. The genetic operators described in subsection 3.3 can be applied λ =10 times at every step of the evolution, with a *σ* = 0.9 strength, and an *α* = 0.9 inertia. To rank protocols by fitness, the DSE engine uses the pair of values *f* = [*f*_0_, *f*_1_] (Equation 1 and Equation 2), measuring precision and coverage of the target, respectively.

### 4.2 Experimental Results

As of the writing of this document, the experiments have been running for 39 days on an Intel(R) Xeon(R) CPU E5-2680 @ 2.70GHz with 64 GB RAM, evaluating 884 protocols along 50 generations.

Results obtained demonstrate that (1) the proposed framework proves capable of automatically generating a protocol for the simulated biofabrication of the illustrated target product and use case and that (2) the DSE can drive the optimization toward protocols with better fitness expressed as similarity to a target product.

Figure 4 (left) shows a 2D view of the Spatial Grid at the beginning (t=0) and end (t=1500) of the simulation of the best protocols identified during different generations of the optimization process. On the right panel of the figure, the chart shows the evolution of the fitness of best protocols along generations. This plot highlights two trends for the fitness of the best protocols consistent with observations performed on all the 884 best protocols (data not shown). Some best protocols (generations 3, 11, 27) exhibit higher precision (*f*_0_), while others (generations 1, 23) higher coverage (*f*_1_). Therefore, the best protocol might be selected from both clusters, depending on the maximum lifetime of the individuals and the advancement of the Pareto front.

**Fig. 4.**
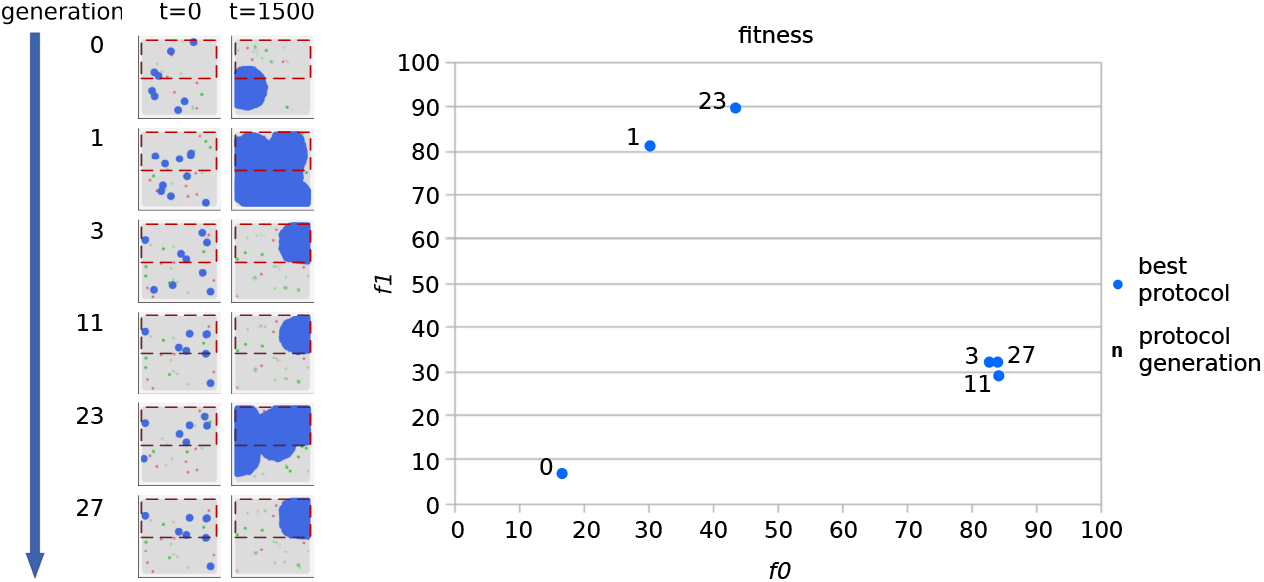
Fitness trend during optimization. Left: product obtained by the best protocols at the beginning and end of the simulation. The dashed red box highlights the target area. Right: fitness of the best protocols.

Figure 5 shows the evolution of the Pareto front, taking into account only the protocols evaluated in the same generation to which the best protocols belong. The best protocol, indicated by a green arrow, is chosen by μGP among the red dots in the image. According to μGP definition of the `MultiObjective` optimization, *“two fitness may be equal, may dominate each other, meaning that all the components of one fitness are greater or equal to the corresponding parts of the other, or they may be not comparable”* [20].

**Fig. 5.**
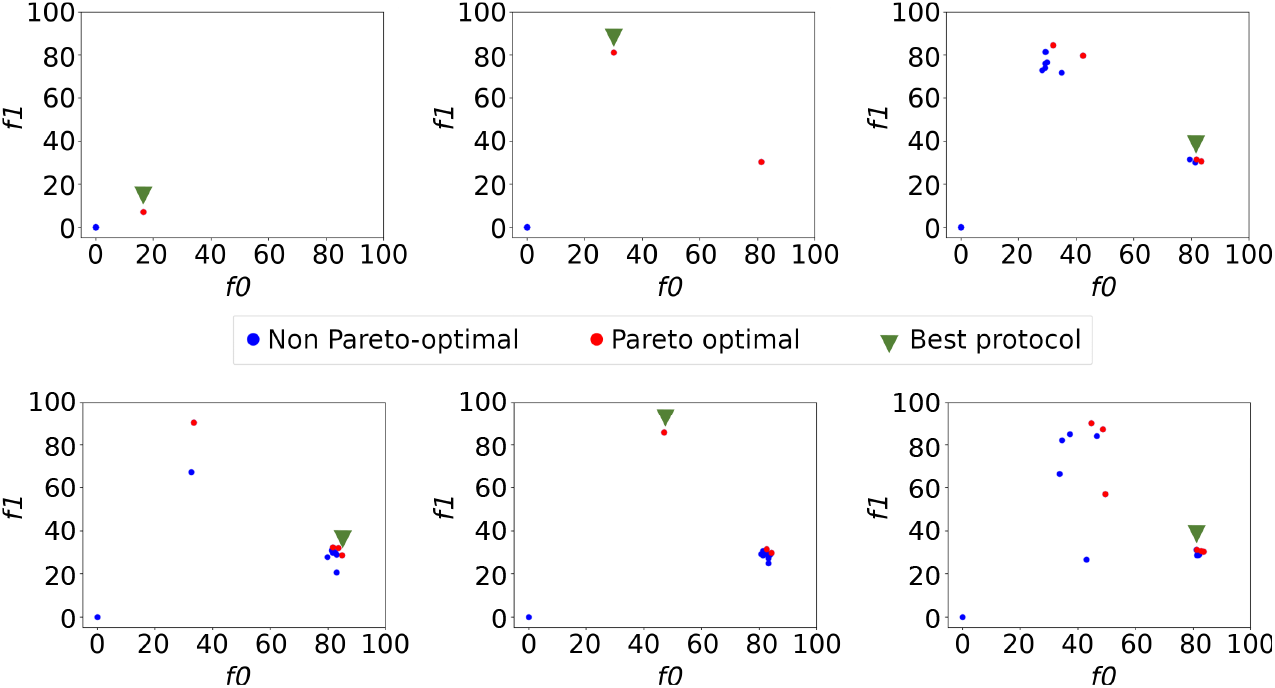
Evolution of the Pareto front. Each chart reports the fitness of the individuals from the same generation, with the Pareto-dominant protocols highlighted in red. The green arrow pinpoints the best individual emerged from that generation. Only generations 0, 1, 3 (top) and 11, 23, 27 (bottom) are shown.

Figure 6 shows per each generation (rows) different initial configurations of the cells (blue), provided by the placing section of the protocols at the beginning of each simulation. 2D views of the Spatial Grid at t=0 represent different simulations for each generation, corresponding to other simulated protocols. At the beginning of the optimization process (generations 0, 1, and 2, top three rows), the DSE engine generates protocols as random individuals. After several rounds of evolution (generations 48 and 49, bottom couple of rows), the initial configuration of cell placing shows more consistency among different simulated protocols.

**Fig. 6.**
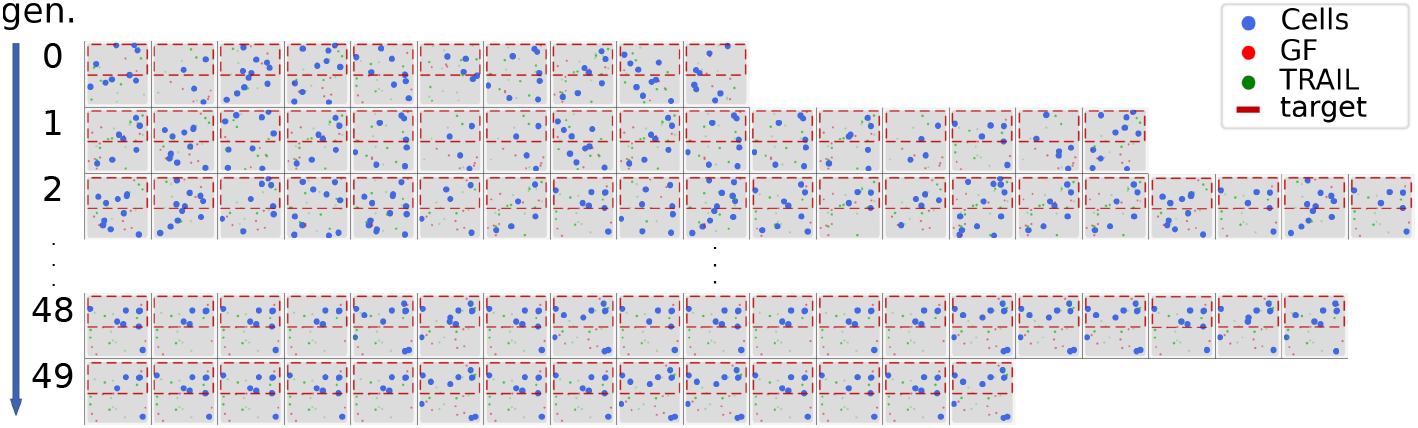
Initial configuration of the cells (blue) obtained from the simulated protocols at the beginning of the simulation. At the beginning of the optimization process, the top three rows are from generations 0–2 (random individuals). The bottom two after several rounds of evolution (generations 48 and 49). The red and green clouds represent Trail and GF signals, respectively. The dashed red box highlights the target area.

Indeed, both Figure 4 and Figure 6 show that, through the generations, the initial placing of the cells gradually shifts and concentrates towards the target area. Figure 7 shows the same simulated protocols at the end of the simulation (t=1500, or t corresponding to death of all the cells). In the first three generations (top three rows), few protocols guarantee cells survival to the end of the simulation. Indeed, those simulations stopped before executing the 1,500 steps (we show the last simulation step with alive cells), corresponding to very low protocol fitness. The two bottom rows (generations 48 and 49) demonstrate that those very poorly fit individuals have permanently given way to protocols that are indeed able to grow conglomerates of cells.

**Fig. 7.**
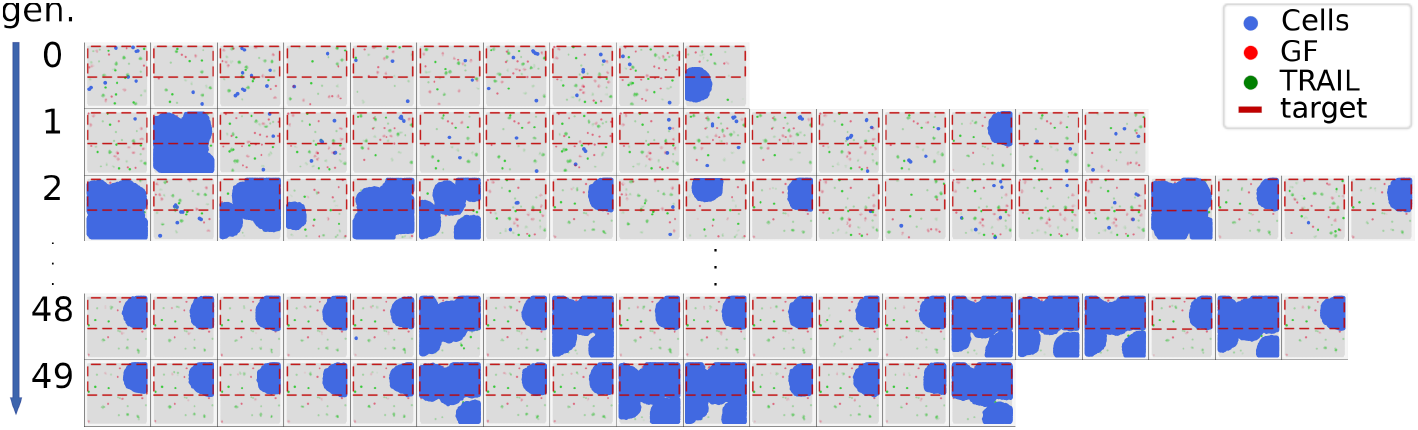
Final configuration of the cells (blue) obtained from the simulated protocols after 1500 simulation steps. At the beginning of the optimization process, the top three rows are from generations 0–2 (random individuals). The bottom two rows (generations 48 and 49) show the progress after several rounds of evolution. The red and green clouds represent Trail and GF signals, respectively. The dashed red box highlights the target area.

Figure 7 indicates the same two-fold tendencies that Figure 4 discovered: some of the fittest protocols yield higher precision, others higher coverage.

The source code and the results obtained are available on GitHub and archived in Zenodo [6].

## 5 Conclusions

In this work, we presented a simulation-optimization methodology for generating biofabrication protocols and a co-simulation framework supporting our strategy. To the best of our knowledge, we are the first to propose this kind of approach. We chose the human epithelium as a use case to validate our methodology and demonstrate the developed framework’s usefulness.

Our results are twofold. First, the prototype framework backing our simulations helps build flexible loosely-coupled co-simulation scenarios. Secondly, the preliminary experimental results show that the proposed approach might provide viable support to biofabrication process design.

In the future, we plan to expand this work in several directions, first, by addressing hyperparameter optimization for the DSE engine. That is the exploration and tuning of optimal GA parameters. Second, by integrating better quantitative models for the realization of more accurate digital twins for both the biofabrication process and the modeled biological system. Finally, we plan to extend our use case by adding cells differentiation so that their diverse functional and phenotypical types let us build more complex products. We are already taking steps towards a massive parallelization, which would allow faster experimentation of larger and more complex biological systems.

While still in its infancy, we can foresee this new methodology as the first step towards standard and automated design in biofabrication.

